# Sensory tuning does not match behavioral relevance of communication signals in free-living weakly electric fish

**DOI:** 10.1101/114249

**Authors:** Jörg Henninger, Rüdiger Krahe, Frank Kirschbaum, Jan Grewe, Jan Benda

**Affiliations:** Institut für Neurobiologie, Eberhard Karls Universität, Auf der Morgenstelle 28E, 72076 Tübingen, Germany; Lebenswissenschaftliche Fakultät, Humboldt-Universität zu Berlin, Philippstr. 13, 10115 Berlin, Germany; McGill University, Department of Biology, 1205 Ave. Docteur Penfield, Montreal, Quebec H3A 1B1, Canada

## Abstract

Sensory systems evolve in the ecological niches each species is occupying. Accordingly, the tuning of sensory neurons is expected to be adapted to the statistics of natural stimuli. For an unbiased quantification of sensory scenes we tracked natural communication behavior of the weakly electric fish *Apteronotus rostratus* in their Neotropical rainforest habitat with high spatio-temporal resolution over several days. In the context of courtship and aggression we observed large quantities of electrocommunication signals. Echo responses and acknowledgment signals clearly demonstrated the behavioral relevance of these signals. The known tuning properties of peripheral electrosensory neurons suggest, however, that they are barely activated by these obviously relevant signals. Frequencies of courtship signals are clearly mismatched with the frequency tuning of neuronal population activity. Our results emphasize the importance of quantifying sensory scenes derived from freely behaving animals in their natural habitats for understanding the evolution and function of neural systems.

## Introduction

Sensory systems evolve in the context of species-specific natural sensory scenes (Lewicki et al., 2014). Consequently, naturalistic stimuli have been crucial for advances in understanding the design and function of neural circuits in sensory systems, in particular the visual (Laughlin, 1981; Olshausen and Field, 1996; Betsch et al., 2004; Gollisch and Meister, 2010) and the auditory system (Theunissen et al., 2000; Smith and Lewicki, 2006; Clemens and Ronacher, 2013). Communication signals are natural stimuli that are, by definition, behaviorally relevant (Wilson, 1975; Endler, 1993; Bradbury and Vehrencamp, 2011). Not surprisingly, acoustic communication signals, for example, have been reported to evoke responses in peripheral auditory neurons that are highly informative about these stimuli (Rieke et al., 1995; Machens et al., 2005). However, other stimulus ensembles that do not strongly drive sensory neurons may also be behaviorally relevant and equally important for understanding the functioning of neural systems. Unfortunately, they are often neglected in electrophysiological studies, because they do not evoke obvious neural responses (Olshausen and Field, 2005).

To avoid this bias, we describe behaviorally relevant sensory scenes that we recorded in an animal’s natural habitat. We then discuss the estimated resulting stimulus properties in the light of known encoding properties of the respective sensory system. Tracking freely behaving and unrestrained animals in natural environments is notoriously challenging (Rodriguez-Munoz et al., 2010). We took advantage of the continuously generated electric organ discharge (EOD; Fig. 1 A) of gymnotiform weakly electric fish, to track their movements and electrocommunication signals without the need of tagging individual fish.

**Figure 1:**
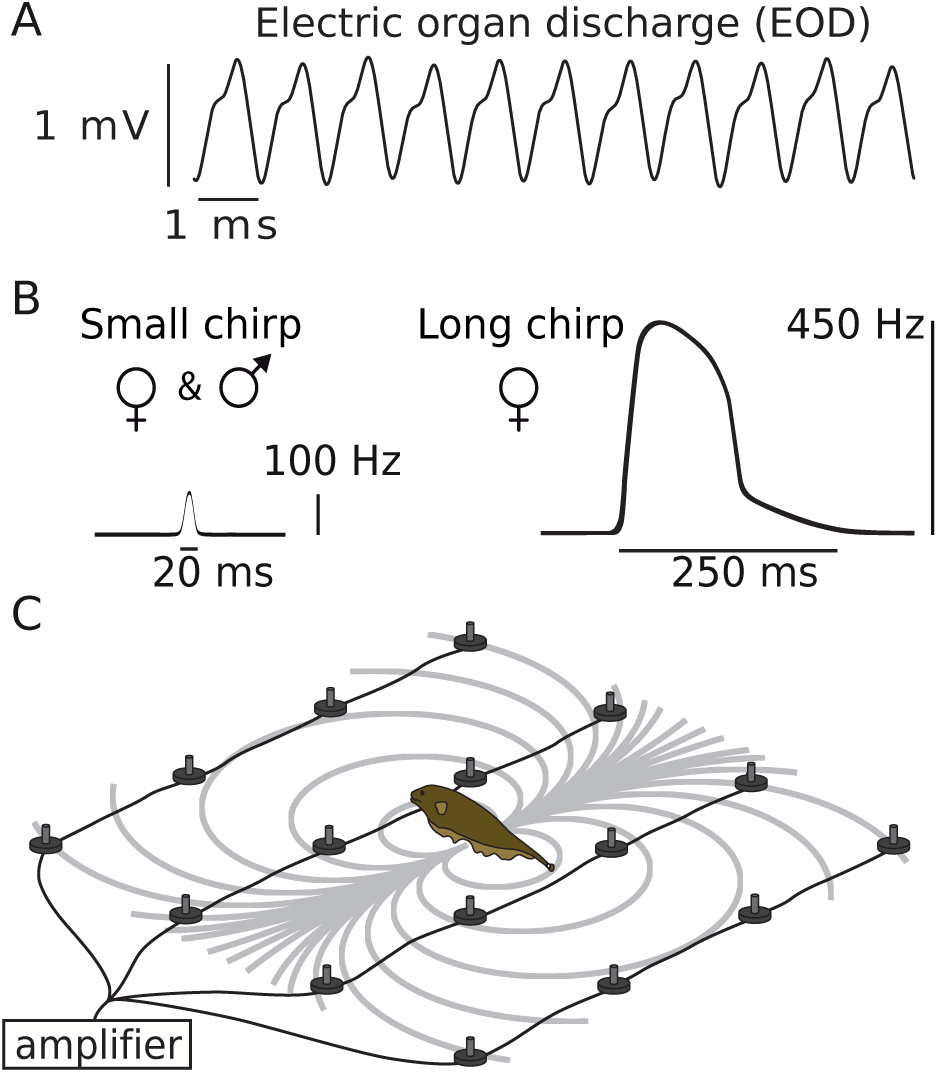
Monitoring electrocommunication behavior in the natural habitat. A) EOD waveform of *A. rostratus*. B) Transient increases of EOD frequency, called small and long chirps, function as communication signals. C) The EOD generates a dipolar electric field (gray isopotential lines) that we recorded with an electrode array, allowing to monitor fish interactions with high temporal and spatial acuity. The following figure supplements are available for figure 1: Figure 1 — figure supplement 1: Field site and the electrode array positioned in a stream.

The quasi-sinusoidal EOD together with an array of electroreceptors distributed over the fish’s skin (Carr et al., 1982) forms an active electrosensory system used for prey capture (Nelson and MacIver, 1999), navigation (Fotowat et al., 2013), and communication (Smith, 2013). Both, the EOD alone and its modulations, function as communication signals that convey information about species, sex, status and intent of individuals: e.g., (Hagedorn and Heiligenberg, 1985; Stamper et al., 2010; Fugère et al., 2011). In *Apteronotus* several types of brief EOD frequency excursions called “chirps”(Fig. 1 B) have been studied extensively in the laboratory (e.g., Engler and Zupanc, 2001) and have been associated with courtship (Hagedorn and Heiligenberg, 1985), aggression (Zakon et al., 2002), and the deterrence of attacks (Hupé and Lewis, 2008). P-unit tuberous electroreceptors encode amplitude modulations of the EOD arising in communication contexts (Bastian, 1981a). Their frequency tuning is crucial for the encoding of amplitude modulations resulting from chirps (Benda et al., 2005; Walz et al., 2014).

We describe, for the first time, electrocommunication behavior of weakly electric fish recorded in their natural neotropical habitat with high temporal and spatial resolution. We found extensive chirping interactions on timescales ranging from tens of milliseconds to minutes in the context of courtship. In a complementary breeding experiment we confirmed the synchronizing role of chirping in spawning. Surprisingly, we found a strong mismatch between properties of courtship signals extracted from our outdoor recordings and the previously described frequency tuning of the respective P-type electroreceptor afferents recorded in electrophysiological experiments. Our data demonstrate that the tuning of receptor neurons can be surprisingly mismatched with respect to the properties of important sensory stimuli, but that sensory systems are nevertheless very well able to process these stimuli.

## Results

We recorded the EODs of weakly electric fish in a stream in the Panamanian rainforest by means of a submerged electrode array at the onset of their reproductive season in May, 2012 (Fig. 1 C, Fig. 1 — figure supplement 1, movie M 1). Individual gymnotiform knifefish, *Apteronotus rostratus*, were identified and their movements tracked continuously based on the species- and individual-specific frequency of their EOD (EOD *f ≈* 580 to 1050 Hz). In these recordings we detected several types of “chirps” emitted during courtship and aggression (Fig. 1 B). This approach allowed us to reconstruct social interactions in detail (Fig. 2, movies M 2 and M 3) and evaluate the associated sensory scenes experienced by these fish in their natural habitat.

**Figure 2:**
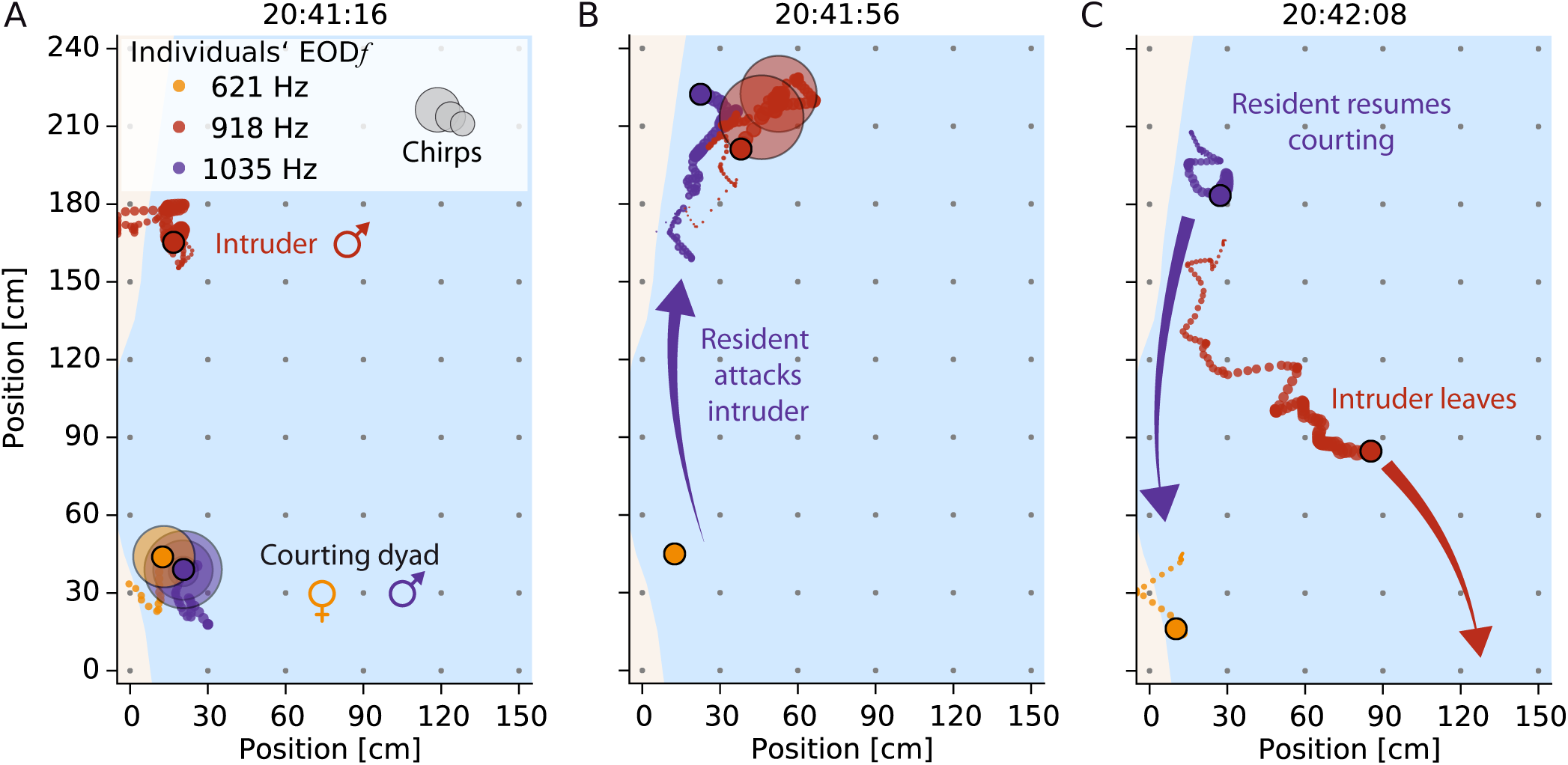
Snapshots of reconstructed interactions of weakly electric fish. See movie M 2 for an animation. The current fish position is marked by filled circles. Trailing dots indicate the positions over the preceding 5 s. Colors label individual fish throughout the manuscript. Large transparent circles denote occurrence of chirps. Gray dots indicate electrode positions, and light blue illustrates the water surface. The direction of water flow is from top to bottom. A) Courting female (orange) and male (purple) are engaged in intense chirping activity. An intruder male (red) lingers at a distance of about one meter. B) The courting male attacks (purple arrow) the intruder who emits a series of chirps and, C) leaves the recording area (red arrow), while the resident male resumes courting (red arrow).

### Electrocommunication in the wild

We focused on two relevant communication situations, i.e., courtship and aggressive dyadic interactions. In total, we detected 54 episodes of short-distance interactions that we interpreted as courtship (see below) between low-frequency females (EOD *f <* 750 Hz, *n*=2) and high-frequency males (EOD *f >* 750 Hz, *n* = 6), occurring in 2 out of 5 nights. Courting was characterized by extensive production of chirps (Fig. 2 A) by both males and females — with up to 8 400 chirps per individual per night (Fig. 3). Most chirps were so-called “small chirps”, characterized by short duration (*<* 20 ms) EOD *f* excursions of less than 150 Hz and minimal reduction in EOD amplitude (Engler and Zupanc, 2001) (Fig. 1 B and Fig. 4). Only females emitted an additional type of chirp in courtship episodes, the “long chirp” (Fig. 1 B and Fig. 4), with a duration of 162 *±* 39 ms (*n* = 54), a large EOD *f* excursion of about 400 Hz, and a strong decrease in EOD amplitude (Hagedorn and Heiligenberg, 1985). Per night and female we observed 9 and 45 long chirps, respectively, generated every 3 to 9 minutes (1st and 3rd quartile), between 7 pm and 1 am (Fig. 3 A). Occasionally, courtship was interrupted by intruding males, leading to aggressive interactions between resident and intruder males (see below).

**Figure 3:**
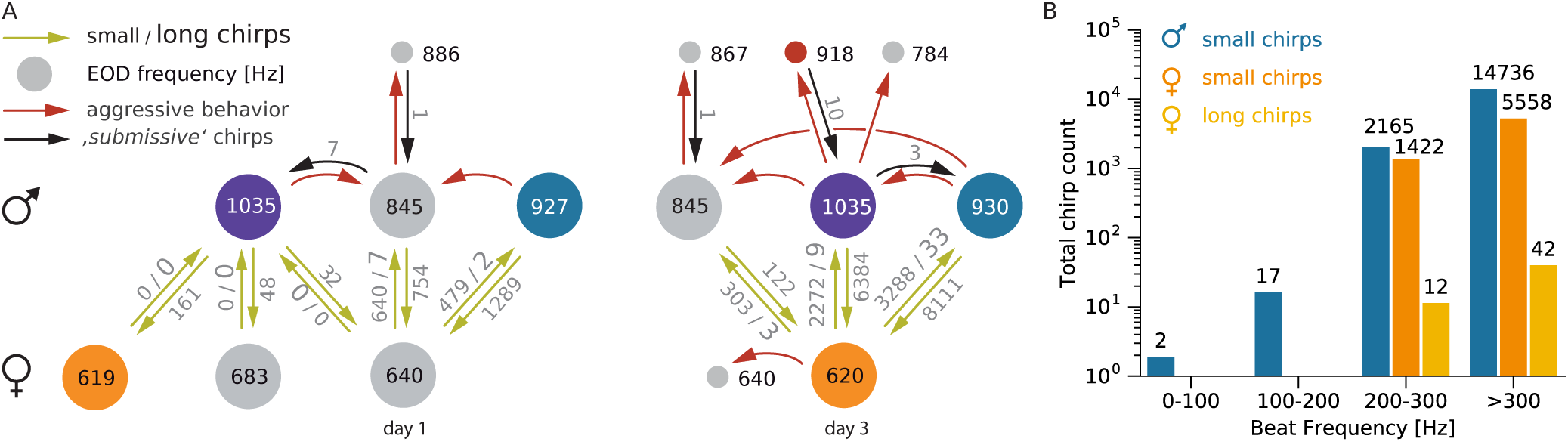
Social interactions and chirping. A) Ethogram of interacting *A. rostratus* individuals (colored circles) displaying their social relationships. The ethogram is based on data from 2012-05-10 (night 1) and 2012-05-12 (night 3) and illustrates the number and EOD frequencies of interacting fish as well as the number of emitted chirps that have been analyzed in this study. The numbers within circles indicate the EOD *f* s of each fish in Hertz. Fish with similar EOD *f* s on day 1 and day 3 may have been the same individuals. Green arrows and associated numbers indicate the numbers of small chirps and long chirps emitted in close proximity (*<* 50 cm). Red arrows indicate aggressive behaviors, and black arrows the number of small chirps emitted during aggressive interactions. B) Histogram of chirp counts as a function of beat frequency (bin-width: 100 Hz). Note logarithmic scale used for chirp counts.

**Figure 4:**
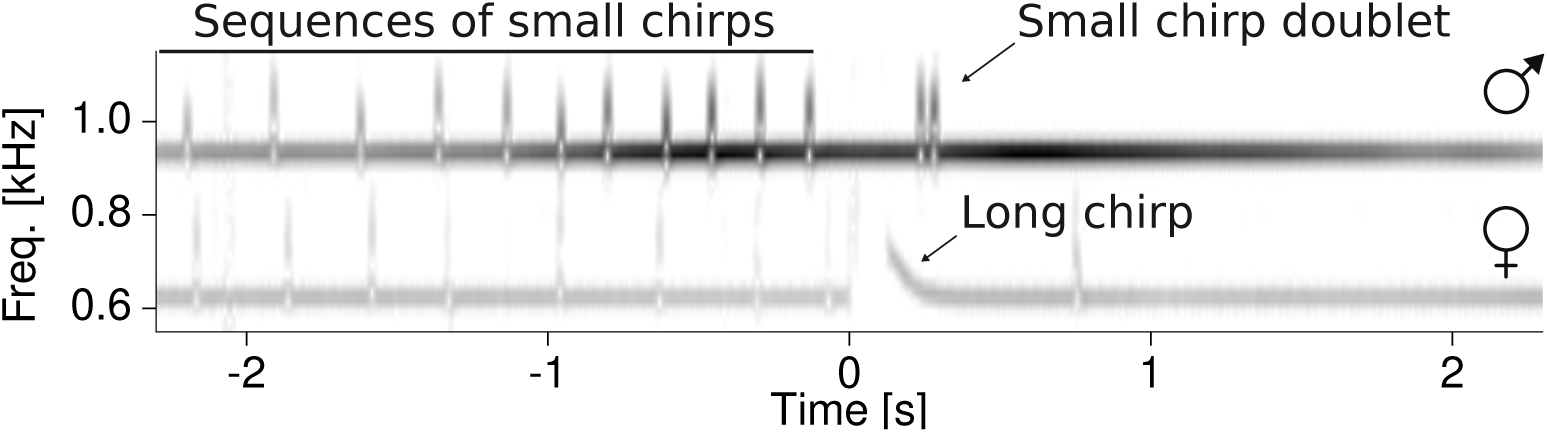
Spectrogram of stereotyped courtship chirping. The example spectrogram (audio A 1) shows EOD *f* s of a female (620 Hz, same as in Fig. 2) and a male (930 Hz) and their stereotyped chirping pattern during courtship: the two fish concurrently produce series of small chirps before the female generates a long chirp. The long chirp is acknowledged by the male with a chirp-doublet that in turn is often followed by one or more small chirps emitted by the female. For statistics see text, Fig. 5, and Fig. 6.

### Courtship chirping

Roaming males approached and extensively courted females by emitting large numbers of small chirps (Fig. 3 A). Courtship communication was highly structured, with female long chirps playing a central role. Long chirps were preceded by persistent emission of small chirps by the male with rates of up to 3 Hz (Figs. 5 A, C). Immediately before the long chirp, the female small-chirp rate tripled from below 1 Hz to about 3 Hz within a few seconds. The male chirp rate followed this increase until the concurrent high-frequency chirping of both fish ceased after the female long chirp. These chirp episodes were characterized by close proximity of the two fish (*<* 30 cm, Fig. 5 B, D). Long chirps were consistently acknowledged by males with a doublet of small chirps (Fig. 4) emitted 229 *±* 31 ms after long chirp onset (*n* = 53 measured in 5 pairs of interacting fish, Fig. 3 A). The two chirps of the doublet were separated by only 46 *±* 6 ms, more than seven-fold shorter than the most prevalent chirp intervals (Fig. 5 — figure supplement 1). Finally, the female often responded with a few more loosely timed small chirps about 670 *±* 0.182 ms after the long chirp (time of first chirp observed in *n* = 33 of the 40 episodes shown in fig. 5). The concurrent increase in chirp rate, its termination by the female long chirp, the male doublet, and the final response by small chirps of the female stood out as a highly stereotyped communication motif that clearly indicates fast interactive communication.

**Figure 5:**
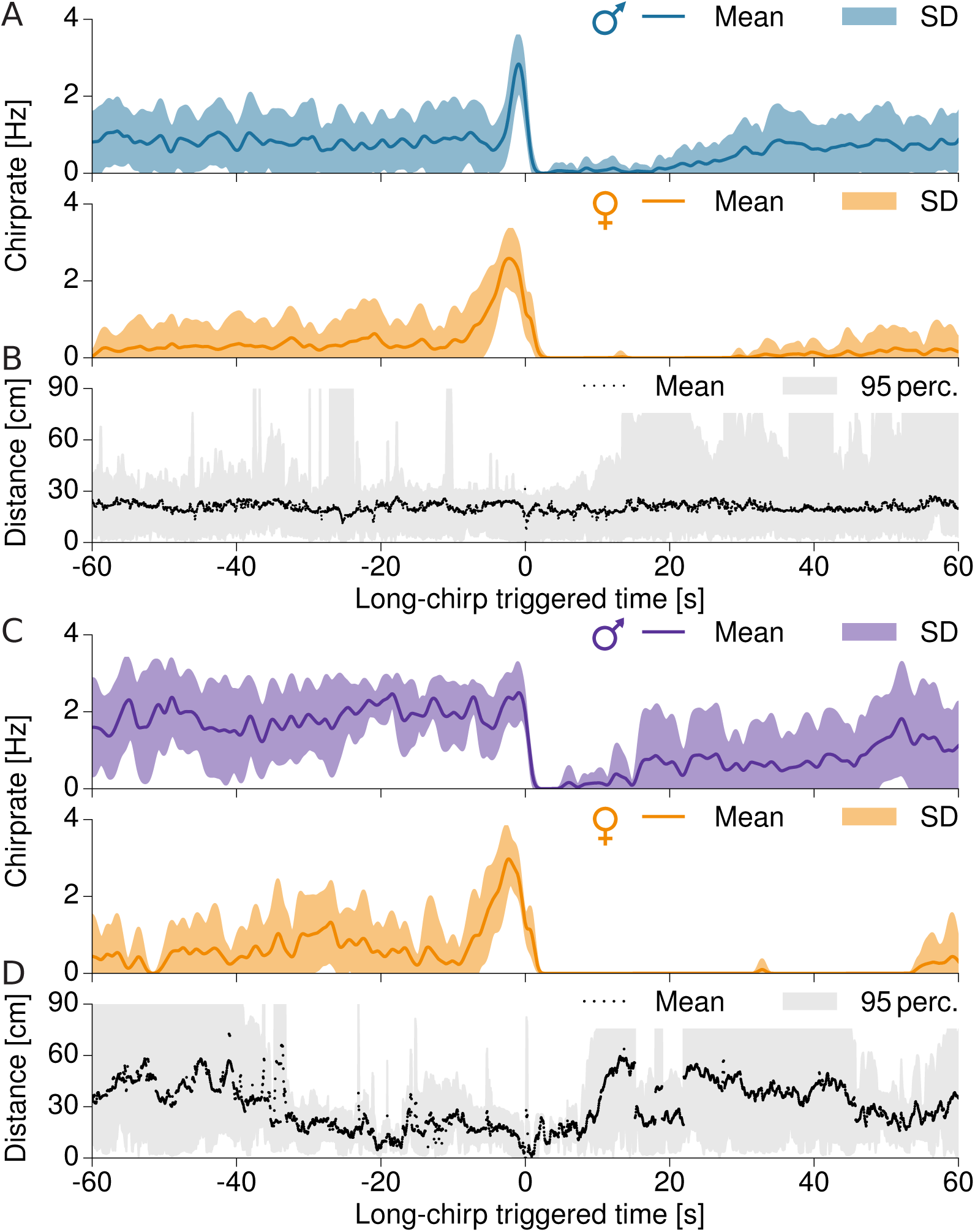
Temporal structure of courtship chirping of two example pairs. A) Average rate of small chirps of a male (top, EOD *f* = 930 Hz) courting a female (bottom, EOD *f* = 620 Hz, *n* = 32 episodes, same pair as in Fig. 4, beat frequency is 310 Hz). B) Corresponding distance between the courting male and female. C, D) Same as in A and B for the pair shown in Fig. 2 (same female as in panel A and B, male EOD *f* = 1035 Hz, beat frequency 415 Hz, *n* = 8 episodes). Time zero marks the female long chirp. Bands mark 95-percentiles. The following figure supplements are available for figure 5: Figure 5 — figure supplement 1: Interchirp-interval distributions of small chirps. Figure 5 — figure supplement 2: Raster plots of small chirps.

### Males echo female chirps

On a sub-second timescale, male chirping was modulated by the timing of female chirps (Figs. 6 A, C). Following a female small chirp, male chirp probability first decreased to a minimum at about 75 ms (significant in 4 out of 5 pairs of fish) and subsequently increased to a peak at about 165 ms (significant in 4 out of 5 pairs of fish). In contrast to males, females did not show any echo response (Figs. 6 B, D) — they timed their chirps independently of the males’ chirps.

**Figure 6:**
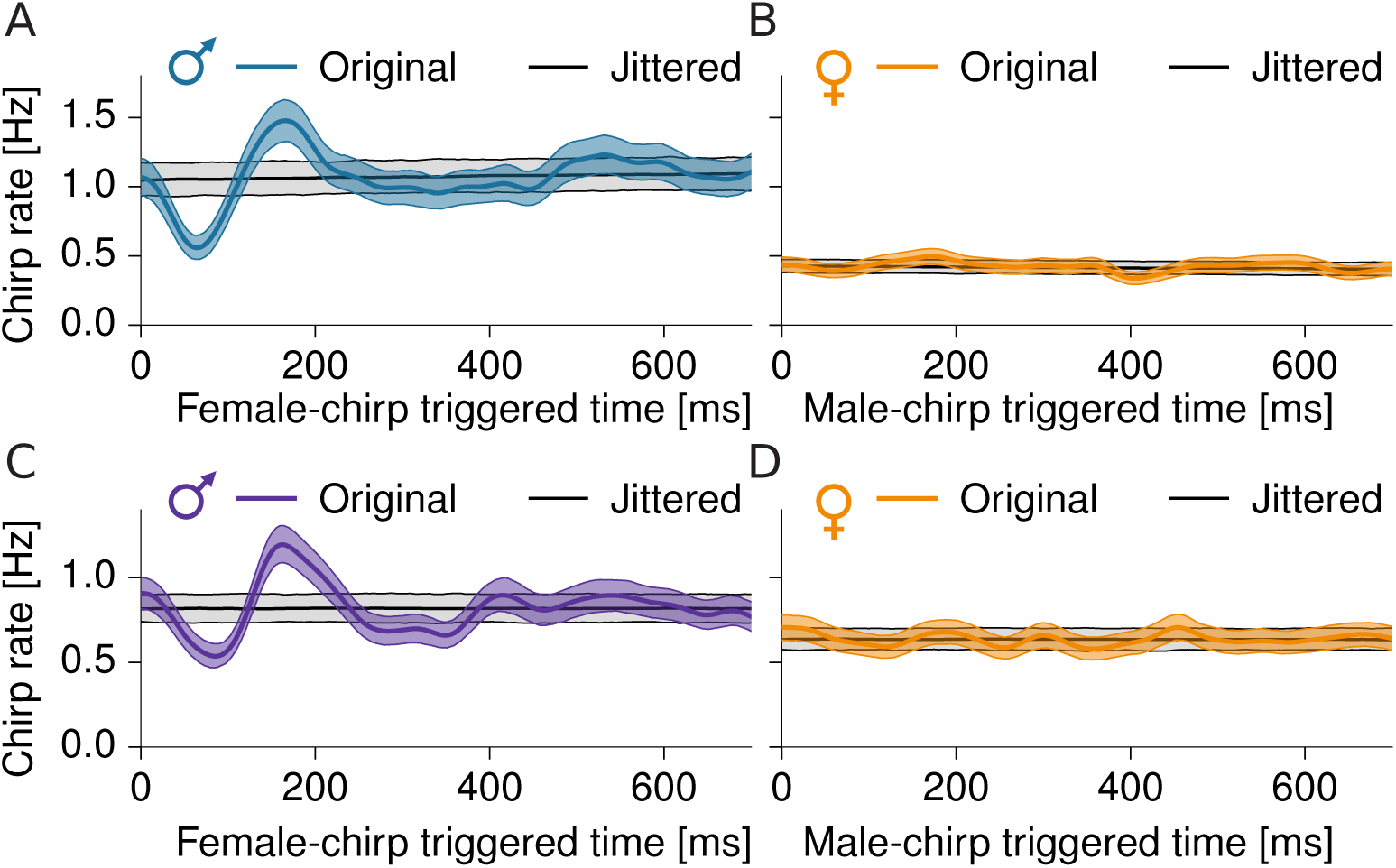
Fine structure of courtship chirping. Shown are cross-correlograms of chirp times, i.e. chirp rate of one fish relative to each chirp of the other fish (median with 95 % confidence interval in color), of the same courting pairs of fish as in Fig. 5. Corresponding chirp rates and confidence intervals from randomly jittered, independent chirp times are shown in gray. A, C) Male chirping is first significantly inhibited immediately after a female chirp (A: at 64 ms, Cohen’s *d* = 9.3, *n* = 2565 female chirps, C: at 85 ms, Cohen’s *d* = 7.1, *n* = 3213 female chirps) and then transiently increased (A: at 166 ms, *d* = 5.9, C: at 162 ms, *d* = 7.5). B, D) Female chirps are timed independently of male chirps (B: maximum *d* = 2.8, *n* = 2648 male chirps, D: maximum *d* = 1.9, *n* = 2178 male chirps).

### Competition between males

A second common type of electro-communication interaction observed in our field data was aggressive encounters between males competing for access to reproductively active females. These aggressive interactions were triggered by intruding males that disrupted courtship of a resident, courting dyad. Resident males detected and often attacked intruders over distances of up to 177 cm, showing a clear onset of directed movement toward the intruder (Fig. 2 C, movie M 2). In 5 out of 12 such situations a few small chirps indistinguishable from those produced during courtship were emitted exclusively by the retreating fish (Fig. 3 A). We observed a single rise, a slow, gradual increase in EOD *f* (Zakon et al., 2002), emitted by a retreating intruder fish.

### Synchronization of spawning

We investigated the role of the female long chirp in a breeding experiment in the laboratory (Kirschbaum and Schugardt, 2002) by continuously recording and videotaping a group of 3 males and 3 females of the closely related species *A. leptorhynchus* (de Santana and Vari, 2013) over more than 5 months. Scanning more than 1.3 million emitted chirps, we found 76 female long chirps embedded in communication episodes closely similar to those observed in *A. rostratus* in the wild (compare Fig. 7 B with Fig. 4). Eggs were only found after nights with long chirps (six nights). The number of eggs found corresponded roughly to the number of observed long chirps, supporting previous anecdotal findings that *Apteronotus* females spawn single eggs during courtship episodes (Hagedorn and Heiligenberg, 1985). The associated video sequences triggered on female long chirps show that before spawning females swim on their side close to the substrate, e.g., a rock or a filter, while the male hovers in the vicinity of the female and emits chirps continuously (movie M 4). In the last seconds before spawning, the female starts to emit a series of chirps, whereupon the male approaches the female. A fraction of a second before the female emits its long chirp, the male pushes the female and retreats almost immediately afterwards (Fig. 7). It seems highly likely that this short episode depicts the synchronized release of egg and sperm.

**Figure 7:**
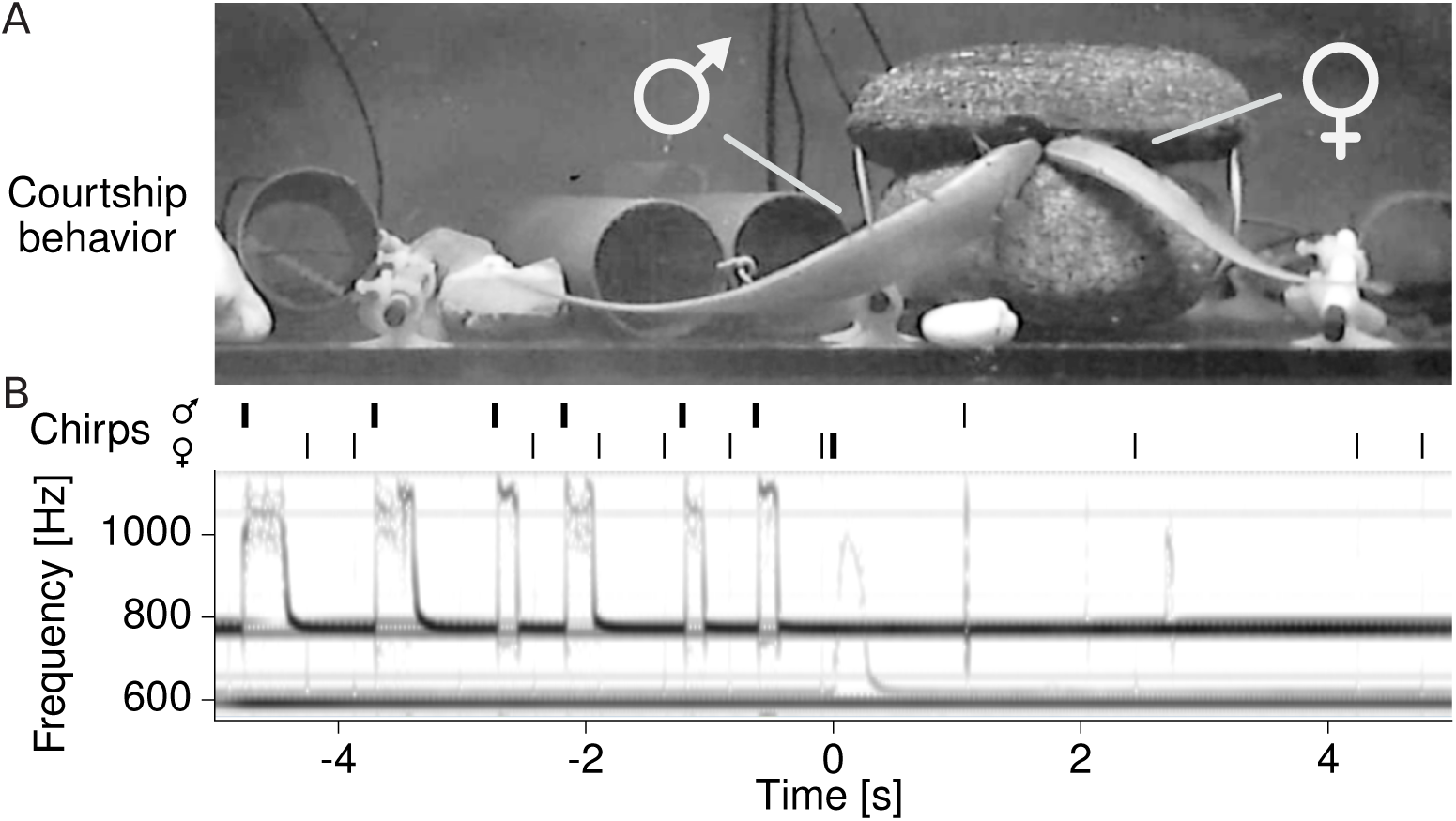
Synchronizing role of the female long chirp in spawning. A) Simultaneous video (snapshot of movie M 4) and B) voltage recordings (spectrogram) of *A. leptorhynchus* in the laboratory demonstrate the synchronizing function of the female long chirp (at time zero; trace with EOD *f* = 608 Hz baseline frequency) in spawning. In contrast to *A. rostratus,* male *A. leptorhynchus* generate an additional, long chirp type before spawning (top trace with EOD *f* = 768 Hz baseline frequency). Chirp onset times of the male and the female are marked by vertical bars above the spectrogram. Thick and thin lines indicate long and short duration chirps, respectively.

### Statistics of natural stimuli

In a final step, we deduced the statistics of natural electrosensory stimuli resulting from the observed communication behaviors to be able to relate it to the known physiological properties of electrosensory neurons in the discussion. Superposition of a fish’s EOD with that of a nearby fish results in a periodic amplitude modulation, a so-called beat. Both frequency and amplitude of the beat provide a crucial signal background for the neural encoding of communication signals (Benda et al., 2005; Marsat et al., 2012; Walz et al., 2014). The beat frequency is given by the difference between the two EOD *f* s and the beat amplitude equals the EOD amplitude of the nearby fish at the position of the receiving fish (Fotowat et al., 2013).

The EOD amplitude and thus the beat amplitude decay with distance. We measured this decay directly from the data recorded with the electrode array (Fig. 8 A). The median EOD field amplitude at 3 cm distance was 2.4 mV*/*cm (total range: 1.4–5.1 mV*/*cm). The electric field decayed with distance according to a power law with exponent 1.28 *±* 0.12 (*n* = 9). This is less than the exponent of 2 expected for a dipole, because the water surface and the bottom of the stream distort the field (Fotowat et al., 2013). Small and long chirps emitted during courtship and small chirps emitted by retreating intruder males occured at small distances of less than 32 cm (Fig. 8 B). In contrast, two behaviors involving intruding males occurred at large distances (Fig. 8 C): (i) Intruding males initially often lingered at distances larger than 70 cm from the courting dyad (*n* = 8, median duration 58.5 s; e.g., Fig. 2 A, movie M 2), consistent with assessment behavior (Arnott and Elwood, 2008). (ii) The distances at which resident males started to attack intruders ranged from 20 cm to 177 cm (81 *±* 44 cm, *n* = 10, Fig. 2 B, movie M 3). At the largest observed attack distance of 177 cm, we estimated the electric field strength to be maximally 0.34 μV*/*cm, assuming the fish were oriented optimally.

**Figure 8:**
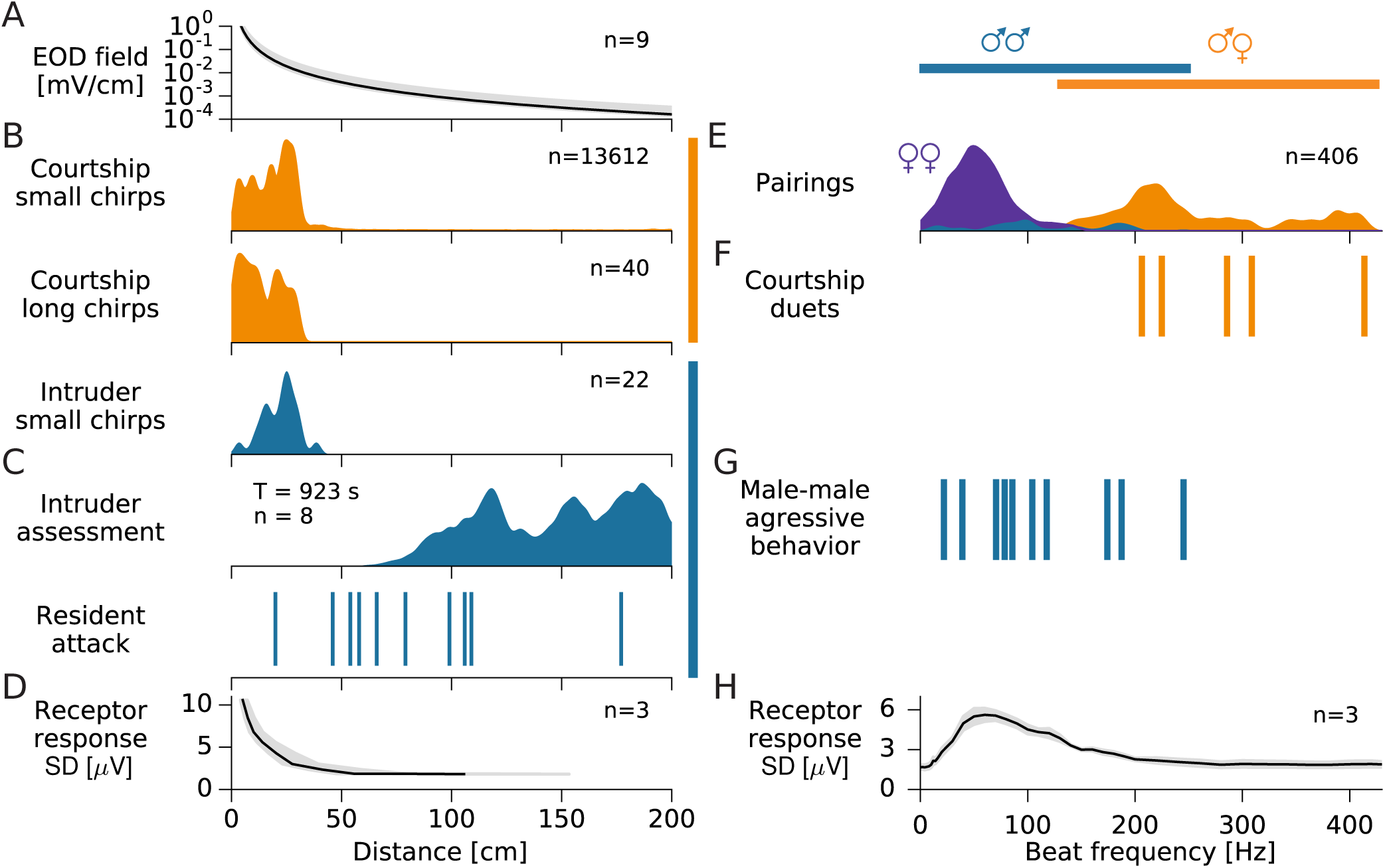
Statistics of behaviorally relevant natural stimuli. A) Maximum electric field strength as a function of distance from the emitting fish (median with total range). B) Small and long chirps in both courtship and aggression contexts are emitted consistently at distances below 32 cm. C) Intruder assessment and initiation of attacks by residents occur at much larger distances (movie M 3). D) Averaged activity of the electroreceptor afferent population rapidly declines with distance between two fish (beat frequency 60 Hz). E) Distribution of beat frequencies of all *A. rostratus* appearing simultaneously in the electrode array. blue: male-male, violet: female-female, orange: male-female (*n* = 406 pairings). F) Courtship behaviors occurred at beat frequencies in the range of 205–415 Hz, far from the receptors’ best frequency. G) Aggressive interactions between males occurred at beat frequencies below 245 Hz, better matching the tuning of the electroreceptor afferents. H) Tuning of electroreceptor afferent activity to beat frequency.

All courtship chirping occurred at high beat frequencies (205–415 Hz for the five pairs where the female emitted long chirps, Fig. 8 F and Fig. 3 B). High beat frequencies were not a rare occurrence as the probability distribution of 406 beat frequencies measured from encounters in 5 nights show (Fig. 8 E). From these the 183 male-female encounters resulted in beat frequencies ranging from 99 to 415 Hz. Same-sex interactions, on the other hand, resulted in low beat frequencies up to 245 Hz (Fig. 8 E). Encounters between females were more frequent than between males (187 female versus 36 male encounters). Female EOD *f* s ranged from 585 to 748 Hz and resulted in observed beat frequencies from 1 to 142 Hz. Beat frequencies of 49 Hz were the most frequent among the females (*n* = 187). Male EOD frequencies, on the other hand, span a much larger range from 776 to 1040 Hz, resulting in a broad and flat distribution of beat frequencies spanning 12 to 245 Hz (peak at 98 Hz, *n* = 36). This includes the range of beat frequencies observed at aggressive male-male interactions (Fig. 8 G).

### Discussion

We recorded movement and electrocommunication signals in a wild population of the weakly electric fish, *Apteronotus rostratus*, in their natural neotropical habitat by means of a submerged array of electrodes. A stereotyped pattern of interactive chirping climaxed in a special long chirp emitted by the female that we identified in a breeding experiment as a synchronizing signal for spawning. The long chirp was reliably acknowledged by the male with a chirp doublet. Interactive chirping leading to a long chirp was characterized by echo responses by the male on a 100 ms time-scale and concurrent increases in chirp rate in dyads on a tens-of-seconds time-scale. Courtship chirping occured at distances below 32 cm and on high beat frequencies of up to 415 Hz. In contrast, aggressive interactions between males like attacks and assessment occur at beat frequencies below about 200 Hz and often at distances larger than half a meter. Retreating males occasionally emit small chirps towards the close-by attacking male.

### Communication in the wild and in the laboratory

Animal communication is defined as the transfer of information by a signal generated by the sender that affects the behavior of the receiving animal (Wilson, 1975). Our observations of male echo responses to female chirps occurring reliably within a few tens of milliseconds (Figs. 6 A, C), precisely timed chirp doublets in response to female long chirps (Figs. 4), immediate behavioral reactions of males to female long chirps (Fig. 7, movie M 4), and females slowly raising their chirp rate in response to male chirping and responding to the male’s chirp doublet (Fig. 5) clearly qualify chirps as communication signals in natural conditions. Laboratory studies have found echo responses on similar (Hupé and Lewis, 2008) or slower timescales of more than about 500 ms (Zupanc et al., 2006; Salgado and Zupanc, 2011) exclusively between males. Small chirps tend to be emitted when fish are less close to each other and correlate with reduced numbers of attacks during staged interactions between pairs of freely moving males (Hupé and Lewis, 2008; Hupé et al., 2008). Therefore, small chirps have been suggested to deter aggressive behavior (Hupé and Lewis, 2008). This is consistent with our observation of a submissive function of male-to-male chirping emitted by an intruder while being driven away by a dominant resident male. The number of chirps generated in these aggressive contexts is, however, much lower (1 to 10, median 3, chirps in 5 of 9 pairings, Fig. 3) compared to encounters staged in the laboratory (about 125 chirps per 5 min trial, Hupé and Lewis, 2008). Our field data do not support a function of chirps as signals of aggression and dominance (Triefenbach and Zakon, 2008). These differences may be due to the specific conditions under which the laboratory experiments were conducted, in particular the restricted space.

In so-called “chirp chamber” experiments, were a fish is restrained in a tube and is stimulated with artificial signals mimicking conspecifics, small chirps were predominantly generated by males at beat frequencies well below about 150 Hz, corresponding to same-sex interactions (Bastian et al., 2001; Engler and Zupanc, 2001). In contrast, in our observations on *A. rostratus* in the field and reproductive *A. leptorhynchus* in the laboratory, both male and female fish were almost exclusively chirping in male-female contexts at beat frequencies above about 200 Hz (Fig. 3 B). Also, the maximum small-chirp rates during courtship exceeded maximum rates from previous laboratory observations (Engler and Zupanc, 2001; Hupé and Lewis, 2008). Last but not least, the total number of chirps we recorded in two nights of courtship activity exceeded the so far published number of chirps recorded under artificial laboratory conditions by an order of magnitude (see Smith (2013) for a review). This large number of chirps allowed us to draw firm conclusions about the function of chirping.

### Electric synchronization of spawning by courtship-specific chirps

Previous laboratory studies already suggested a function of the EOD and its modulations in courtship in *A. rostratus* (Hagedorn and Heiligenberg, 1985; Hagedorn, 1988) and in the synchronization of spawning and external fertilization in a pulse-type gymnotiform fish (Silva et al., 2008). Our results provide strong evidence that female long chirps are an exclusive communication signal for the synchronization of egg spawning and sperm release: (i) The female long chirp was the central part of a highly stereotyped duet-like communication pattern between a male and female fish (Fig. 4 and Fig. 5). Multimodal signals synchronizing egg and sperm release have been observed in other aquatic animals (Lobel, 1992; Ladich, 2007). (ii) Fertilized eggs were found at the locations of male-female interaction, and only when the female had produced long chirps in the preceding night. (iii) The period immediately before the female long chirp was characterized by extensive chirp production by the male (Fig. 5), consistent with male courting behavior (Bradbury and Vehrencamp, 2011). (iv) Video sequences triggered on female long chirps clearly demonstrated the special role of the female long chirp (Fig. 7, movie M 4). We hypothesize that the synchronization of external fertilization via extensive electrocommunication gives females good control over who they reproduce with.

The males appear to compete for access to females. This and the observed serial consortship suggest a scramble competition mating system as it is common for animals living in a three-dimensional environment, e.g., fish and birds (Bradbury and Vehrencamp, 2011).

### Encoding of communication signals

Male echo responses to female chirps occurring reliably within a few tens of milliseconds (Figs. 6 A, C), precisely timed chirp doublets (Figs. 4), and long-range assessment and attacks (Fig. 8 C) demonstrate that the respective electrocommunication signals are successfully and robustly evaluated by the electrosensory system, as it is expected for communication signals (Wilson, 1975; Endler, 1993). The electrosensory signals arising in these interactions are dominated by beats, i.e. amplitude modulations arising form the interference of the individual electric fields. Beats and their modulations by chirps are characterized by their amplitude and frequency. How are these signals encoded by the electrosensory periphery?

Gymnotiform weakly electric fish have three types of electroreceptor cells: (i) Ampullary receptors are tuned to low-frequency (*<* 50 Hz) external electric fields and therefore do not encode the EOD with its EOD *f* of more than about 600 Hz (Grewe et al., 2017). They also do not encode chirps in *Apteronotus*, which have no low-frequency component, in contrast to different types of chirps emitted by *Eigenmannia virescens* (Metzner and Heiligenberg, 1991; Stöckl et al., 2014). (ii) T-unit tuberous receptors help to disambiguate the sign of the beat frequency in the context of the jamming avoidance response (JAR) in *Eigenmannia* for beat frequencies below about 20 Hz (Bullock et al., 1972; Rose and Heiligenberg, 1985). They are also able to encode low-frequency amplitude modulations (Carlson and Kawasaki, 2006). Whether and how T-units are able to encode beats with frequencies higher than 20 Hz is, however, not known yet. (iii) P-units, the second and dominant type of tuberous receptor in *Apteronotus* (Carr et al., 1982), encode amplitude modulations of the fish’s EOD in their firing rate (Scheich et al., 1973; Bastian, 1981a; Nelson et al., 1997; Benda et al., 2005; Walz et al., 2014). Thus, P-units are potentially the only type of electroreceptors that are able to encode the wide range of amplitude modulations generated by beats and chirps in *Apteronotus*.

For illustrating the known tuning properties of the P-unit population in Fig. 8 D, H (Bastian, 1981a; Nelson et al., 1997; Benda et al., 2006; Walz et al., 2014), we estimated the population activity of P-unit afferents in *A. leptorhynchus* from the standard deviation of the summed nerve activity. This nerve activity is known to closely match the average tuning of the firing rate of single nerve fibers (Benda et al., 2006; Walz et al., 2014). Note also, that the frequency tuning of P-unit afferents has, to our knowledge, so far never been measured beyond 300 Hz and amplitude tuning never below about 10 μV*/*cm (Bastian, 1981a). In the following we first discuss the relation of the P-unit’s average firing-rate tuning in relation to the natural stimulus statistics we measured in the field. This is then followed by a more detailed discussion on alternative coding strategies.

### Frequency mismatch

P-unit afferents are tuned to beat frequency and have on average strongest firing-rate responses around 60 Hz (Bastian, 1981a; Walz et al., 2014) (Fig. 8 H), covering well the beat frequencies arising from same-sex interactions (Fig. 8 G). Remarkably, all courtship chirping occurred at much higher beat frequencies (205–415 Hz, Fig. 8 F and Fig. 3 B). Even though the beat amplitudes (Fig. 8 A) during these interactions are large at the observed small distances of less than 32 cm (Fig. 8 B), such high-frequency stimuli should evoke only weak P-unit activity. High beat frequencies are not a rare occurrence as the probability distribution of 406 beat frequencies measured from all encounters observed in 5 nights show (Fig. 8 E), demonstrating a clear frequency mismatch between an important group of behaviorally relevant sensory signals and electroreceptor tuning.

### Evolutionary origin of frequency mismatch

This profound mismatch between the tuning of receptor neurons and the high beat frequencies occurring during courtship is unexpected given the many examples of frequency-matched courtship signals in other sensory systems (Rieke et al., 1995; Machens et al., 2005; Kostarakos et al., 2009; Schrode and Bee, 2015). The observed frequency-mismatch is also unexpected from the perspective of the design of animal communication systems (Endler, 1993; Bradbury and Vehrencamp, 2011) and points to selective forces beyond the need for the male to provide strong stimulation to the female sensory system.

The high beat-frequencies in male-female interactions are a consequence of a sexual dimorphism in EOD *f* where males have higher frequencies (here 907 *±* 77 Hz, *n* = 10, Fig. 3 A) than females (here 640 *±* 23 Hz, *n* = 5). In the genus *Apteronotus* the presence, magnitude, and direction of an EOD *f* dimorphism varies considerably across species and thus is evolutionarily labile (Smith, 2013). This points to very specific and subtle selection pressures generating large EOD *f* dimorphisms that offset the reduced sensitivity for mates.

A possible selection pressure generating the dimporphism in EOD *f* could be that females prefer males with the highest EOD frequency that nevertheless are able to generate an echo response (Figs. 6). Since detection of female chirps becomes more difficult with higher beat frequencies (see also below) echoing will also be more difficult. The echo-response then would be an honest signal of the male’s ability to detect female chirps, which is important for a successful fertilization of the eggs. However, as explained below, for females it is even harder to detect chirps on high-frequency beats than for males.

In addition to the substantial metabolic cost of the active electrosensory system incurred by continuously generating an EOD (Salazar et al., 2013; Markham et al., 2016), the frequency mismatch potentially also imposes a computational cost on the sensory system.

### Encoding of high-frequency courtship signals

The many behaviors we observed in our study clearly demonstrate that the fish are able to perceive high-frequency beats and small and long chirps generated on these beats. How are these signals encoded by P-unit electroreceptor afferents despite the frequency mismatch?

The firing-rate tuning of P-units to beat frequencies has been characterized up to 300 Hz by single-unit recordings in both *A. albifrons* and *A. leptorhynchus* (Bastian, 1981a; Nelson et al., 1997; Benda et al., 2006; Walz et al., 2014). All studies agree that the average P-unit response is strongest at beat frequencies of about 30 to 130 Hz and declines almost back to baseline levels at 300 Hz (Fig. 8 H). However, P-units are heterogeneous in their baseline activity (Gussin et al., 2007; Savard et al., 2011; Grewe et al., 2017) and P-units with high baseline rates are likely to exhibit a frequency tuning that extends to higher frequencies than the average tuning of the population (Knight, 1972). This is supported by noticeable stimulus-response coherences that have been measured with narrowband noise stimuli up to 400 Hz (Savard et al., 2011). Most of these studies used rather strong beat amplitudes of more than 10 % of the EOD amplitude. We observed chirp interactions at distances up to 32 cm, corresponding to beat amplitudes from 100 % down to about 1 % of the EOD amplitude (Fig. 8 A). Smaller beat amplitudes result in down-scaled frequency tuning curves (Bastian, 1981a; Benda et al., 2006). In particular, for amplitudes below 10 %, population responses to beat frequencies larger than 200 Hz are close to baseline activity (Benda et al., 2006).

Encoding of small chirps can be understood based on the frequency tuning of P-units (Walz et al., 2014). A chirp transiently increases the beat frequency. The firing-rate response to the chirp differs only as much from the response to the background beat as the frequency response to the chirp differs from the one to the beat. Therefore, we expect chirp coding to be additionally impaired by the low slope of the P-unit’s firing-rate tuning curve at high beat frequencies. Note that there is an asymmetry in chirp encoding: Chirps emitted by females result in more distinct responses in males with their higher EOD *f*, because in this scenario the chirp reduces the beat frequency. On the other hand, a male chirp results in a much weaker activation of the female’s receptor population, because the chirp increases the beat-frequency even more (Walz et al., 2014). This matches the observed asymmetry of the echo response, but other behaviors suggest that the females do perceive the male chirps (Fig. 5).

In contrast to small chirps, long chirps show a strong decrease in EOD amplitude during the chirp. We expect the encoding of long chirps to be based on this amplitude decrease as has been shown for the shorter type-1 chirps (Benda et al., 2006): For the duration of the type-1 chirp the electroreceptor activity reverts back to baseline activity. Encoding of long chirps should therefore not be impaired by the shallow slope of the firing-rate tuning curve, but should only rely on some activity induced by the background beat.

### Encoding of low amplitude beats

Two behaviors involving intruding males occurred within the P-units’ best-frequency range (Fig. 8 G), but at large distances (Fig. 8 C): (i) Intruding males initially often lingered at distances larger than 70 cm from the courting dyad (8 of 16 scenes, median duration 58.5 s; e.g., Fig. 2 A, movie M 2), consistent with assessment behavior (Arnott and Elwood, 2008). The average P-unit population response quickly drops to low signal-to-noise ratios at amplitudes corresponding to inter-fish distances larger than about 50 cm at 60 Hz beat frequency (Fig. 8 D, Bastian, 1981a). Consequently, the signals occurring during assessment activate the P-unit population only weakly. (ii) The distances at which resident males started to attack intruders ranged from 20 cm to 177 cm (81 *±* 44 cm, *n* = 10, Fig. 2 B, movie M 3). At the largest observed attack distance of 177 cm, the electric field strength was estimated to be maximally 0.34 μV*/*cm (assuming the fish were oriented optimally) — a value close to minimum behavioral threshold values of about 0.1 μV*/*cm measured in the laboratory at the fish’s best frequency (Knudsen, 1974; Bullock et al., 1972). Both situations, opponent assessment and decision to attack, therefore evoke weak activity of P-units close to the fish’s perception threshold.

Note that large distances and the associated weak receptor activation occur because of physical necessities. This is in contrast to the high beat frequencies that are the result of an evolutionary adaptation as discussed above.

### Decoding

The robust behavioral responses suggest that the weak activation of electroreceptor neurons is compensated for at later processing stages. The compensating mechanisms likely involve synchrony detection (Middleton et al., 2009; Grewe et al., 2017), averaging (Maler, 2009b; Jung et al., 2016), and feedback connections from higher brain areas, including the telencephalon (Giassi et al., 2012; Krahe and Maler, 2014), that modulate the first stage of electrosensory processing in the hindbrain (Bastian, 1986). P-units converge onto pyramidal cells in the electrosensory lateral line lobe (ELL) (Heiligenberg and Dye, 1982; Bastian, 1981b; Maler, 2009a). The pyramidal cells processing communication signals (Metzner and Juranek, 1997; Krahe et al., 2008; Marsat et al., 2009; Vonderschen and Chacron, 2011; Marsat et al., 2012; Metzen et al., 2016) integrate over 1000 P-units each (Maler, 2009a), their rate tuning curves peak at frequencies similar to or lower than those of P-units (Bastian, 1981b), and their stimulus-response coherences peak well below 100 Hz, but have only been measured up to 120 Hz (Chacron et al., 2003; Chacron, 2006; Krahe et al., 2008). Interestingly, peak coherences are higher in *A. leptorhynchus* than in *Apteronotus albifrons*, pointing to species-specific adaptations (Martinez et al., 2016). Common input from overlapping receptive fields (Maler, 2009a) results in significant spike-time correlations between nearby pyramidal cells at high stimulus amplitudes (Chacron and Bastian, 2008). We expect these correlations to vanish at low stimulus amplitudes such that averaging over pyramidal cells could improve information transmission, as suggested by Jung et al. (2016).

Coding of small chirps by pyramidal cells in the ELL and in the next step of electrosensory processing, the Torus semicircularis, has so far only been studied at beat frequencies below 60 Hz (Marsat et al., 2009; Marsat and Maler, 2010; Vonderschen and Chacron, 2011; Marsat et al., 2012; Metzen et al., 2016). Thus, most electrophysiological recordings from the electrosensory system have been biased to low beat frequencies and strong stimulus amplitudes evoking obvious neuronal responses, overlooking the behaviorally most relevant stimuli. For the visual system a preference of experimenters for responsive neurons has been identified as a major bias in sampling neurons (Olshausen and Field, 2005).

Coding of type-1 chirps in ELL, and Torus, on the other hand, has been measured on the relevant beat frequencies of several hundred Hertz (Benda et al., 2006; Marsat and Maler, 2010). Indeed, Torus neurons have been found that respond relatively selectively to type-1 chirps (Vonderschen and Chacron, 2011; Marsat et al., 2012). We expect similar responses in ELL and Torus for long chirps, because their large increase in EOD frequency and their decrease of EOD amplitude is similar to type-1 chirps. Note, however, that long chirps are about ten times longer in duration than type-1 chirps (Bastian et al., 2001; Zupanc et al., 2006). This demonstrates that neural responses at higher processing levels get more specific and distinct to behaviorally relevant signals. Our data on natural behavior, however, point to further relevant communication signals whose neural representations have not been studied yet.

### Hormonal influences

Depending on the state of the animal, electrosensory tuning may be modified, e.g., by hormones and/or neuromodulators. Steroids are known to modulate EOD *f* and tuning of P-units to EOD *f* (Meyer and Zakon, 1982; Meyer et al., 1987; Dunlap et al., 1998). Serotonin is a neuromodulator that is released in the ELL in response to communication signals (Fotowat et al., 2016) and potentially enhances activity levels in ELL (Deemyad et al., 2013). Activation of muscarinic receptors by acetylcholine has been shown to improve low-frequency coding of pyramidal cells (Ellis et al., 2007). Whether and how hormones or neuromodulators influence coding of high-frequency beats is not known yet.

### Non-optimal coding?

Both large-distance interactions between males and courtship interactions between males and females result in sensory stimuli at the limits of the respective tuning curves (Fig. 8). Depending on the neural code, be it the population rate or the stimulus-response coherence for example, the respective tuning curves vanish sooner or later. We expect any code to eventually vanish at high distances or high frequencies, because of basic physical constraints in the transduction of small stimulus amplitudes (Fig. 8, Bialek, 1987) or in the frequency tuning of fundamental spike-generators (Knight, 1972; Fourcaud-Trocmé et al., 2003). Also the synchrony code suggested by Middleton et al. (2009), that can be observed only in the limit of small stimulus amplitudes (Sharafi et al., 2013; Kruscha and Lindner, 2016; Grewe et al., 2017), vanishes at even smaller amplitudes. Thus we assume that at large distances and high frequencies the response of any code would approach zero despite finite probabilities of the corresponding stimuli that we measured (Fig. 8 C,F). This would violate maximization of mutual information between stimulus and response (Laughlin, 1981).

In this study we quantified the distribution of beat frequencies as they occurred in the mating season (Fig. 8 E). These frequencies occur independently of whether the fish actually engage in courtship or aggressive interactions. In fact, these frequencies are always present, as soon as another fish is close by, i.e. within less than about 2 m. Outside the breeding season the EOD frequencies and thus the resulting beat frequencies may change (Hagedorn and Heiligenberg, 1985). In addition to beats and chirps, amplitude modulations caused by nearby objects, including prey items, are also relevant stimuli (Nelson and MacIver, 1999; Fotowat et al., 2013). These amplitude modulations are low in frequency (*<* 20 Hz) and in case of prey items also low in amplitude (Benda et al., 2013). Interestingly, the firing-rate response to these signals is also low (Nelson et al., 1997), because of a strong spike-frequency adaptation (Benda et al., 2005). Negative interspike-interval correlations (Ratnam and Nelson, 2000) reduce the noise level in this regime, resulting in still high stimulus-response coherences (Chacron et al., 2005) that are conveyed to the pyramidal cells via matched short-term synaptic dynamics (Nesse et al., 2010). Navigation stimuli need to be processed all the time, as are beat frequencies of conspecifics. Thus, the frequency distribution in Fig. 8 E needs to be complemented by another peak at very low stimulus frequencies of about the same height as the distribution of beat frequencies. However, factoring in behavioral relevance, it could well be that outside the breading season the beat frequencies are not relevant any more, although they still could be present. This would weigh the beat-frequencies much less in comparison to electrolocation signals and consequently the observed frequency mismatch would much less violate an optimal infomax code in this scenario.

### Conclusion

Our observations regarding sex-specificity, numbers, and functions of chirps differ substantially from laboratory studies. Limited space and artificial settings may have biased interactions towards aggressive behaviors. In the wild, these aggressive encounters were rare and were accompanied by only little or no chirping. In contrast, courtship scenes stood out as highly structured and long-lasting sequences of high-frequency chirping. We have shown that similar courtship behavior can be reproduced in the laboratory by giving the fish enough time to interact — several months instead of minutes or hours.

The fish robustly responded to courtship signals although courtship signals activate P-unit electroreceptors only weakly at the tail of their tuning curve. The extent of this mismatch in frequency tuning was unexpected given previous, mainly laboratory-based findings (Smith, 2013; Walz et al., 2013). For the first time we observed long-distance interactions between competing males that also emphasize the ability of the electrosensory system to process relevant signals close to threshold reliably. Our field data thus identify important — but so far neglected — stimulus regimes of the electrosensory system and provide further evidence for the existence of sensitive neural mechanisms for the detection of such difficult sensory signals (Gao and Ganguli, 2015).

The analysis of field data from the natural environments a specific species evolved in could point to behaviorally relevant sensory scenes that are otherwise neglected because they do not obviously excite sensory neurons (Olshausen and Field, 2005). Here, this is exemplified for weakly electric fish. For other organisms and sensory sytems field data may as well reveal unexpected sensory scenes. Such difficult and complex signals that nevertheless are behaviorally relevant open new windows for investigating the real challenges faced and solved by sensory systems.

## Materials and methods

### Field site

The field site is located in the Tuira River basin, Province of Darién, Republic of Panamá (fig. 1 — figure supplement 1 A), at Quebrada La Hoya, a narrow and slow-flowing creek supplying the Chucunaque River. Data were recorded about 2 km from the Emberá community of Peña Bijagual and about 5 km upstream of the stream’s mouth (8°15^*I*^13.50^*II*^N, 77°42^*I*^49.40^*II*^W). The water of the creek is clear, but becomes turbid for several hours after heavy rainfall. The creek flows through a moist secondary tropical lowland forest, which, according to local residents, gets partially flooded on a regular basis during the wet season (May – November). The water levels of the creek typically range from 20 – 130 cm at different locations, but can rise temporarily to over 200 cm after heavy rainfall. At our recording site (fig. 1 — figure supplement 1 B), the water level ranged from 20 – 70 cm. The banks of the creek are typically steep and excavated, consisting mostly of root masses of large trees. The water temperature varied between 25 and 27°C on a daily basis and water conductivity was stable at 150 – 160 μS*/*cm. At this field site we recorded four species of weakly electric fish, the pulse-type fish *Brachyhypopomus occidentalis* (about 30 – 100 Hz pulses per second), the wave-type species *Sternopygus dariensis* (EOD *f* at about 40 – 220 Hz), *Eigenmannia humboldtii* (200 – 580 Hz), and *Apteronotus rostratus* (580 – 1100 Hz). We here focused exclusively on *A. rostratus*, a member of the *A. leptorhynchus* species group (brown ghost knifefish (de Santana and Vari, 2013)) and its intraspecies interactions.

### Field monitoring system

Our recording system (Fig. 1 C, fig. 1 — figure supplement 1 B) consisted of a custom-built 64-channel electrode and amplifier system (npi electronics GmbH, Tamm, Germany) running on 12 V car batteries. Electrodes were low-noise headstages encased in epoxy resin (1*×* gain, 10 *×* 5 *×* 5 mm). Signals detected by the headstages were fed into the main amplifier (100*×* gain, 1st order high-pass filter 100 Hz, low-pass 10 kHz) and digitized with 20 kHz per channel with 16-bit amplitude resolution using a custom-built low-power-consumption computer with two digital-analog converter cards (PCI-6259, National Instruments, Austin, Texas, USA). Recordings were controlled with custom software written in C++ (https://github.com/bendalab/fishgrid) that also saved data to hard disk for offline analysis (exceeding 400 GB of uncompressed data per day). Raw signals and power spectra were monitored online to ensure the quality of the recordings. We used a minimum of 54 electrodes, arranged in an 9 *×* 6 array covering an area of 240 *×* 150 cm (30 cm spacing). The electrodes were mounted on a rigid frame (thermoplast 4 *×* 4 cm profiles, 60 % polyamid, 40% fiberglass; Technoform Kunststoffprofile GmbH, Lohfelden, Germany), which was submerged into the stream and fixed in height 30 cm below the water level. Care was taken to position part of the electrode array below the undercut banks of the stream in order to capture the EODs of fish hiding in the root masses. The recording area covered about half of the width of the stream and the hiding places of several electric fish. The maximum uninterrupted recording time was limited to 14 hours, determined by the capacity of the car batteries (2 *×* 70 Ah) and the power consumption of the computer (22 W) and amplifier system (25 W).

### Data analysis

All data analysis was performed in Python 2.7 (www.python.org, https://www.scipy.org/). Scripts and raw data (Panamá field data: 2.0 TB, Berlin breeding experiment: 3.7 TB of EOD recordings and 11.4 TB video files) are available on request, data of the extracted EOD frequencies, position estimates and chirps are available at https://web.gin.g-node.org/bendalab, and some of the core algorithms are accessible at https://github.com/bendalab/thunderfish. Summary data are expressed as means *±* standard deviation, unless indicated otherwise.

### Spectrograms

Spectrograms in Fig. 4 and Fig. 7 B were calculated from data sampled at 20 kHz in windows of 1024 and 2048 data points, corresponding to 51.2 ms and 102.4 ms, respectively, applying a Blackman window function. Sequential windows were shifted by 50 data points (2.5 ms). The resulting spectrograms were interpolated in the frequency dimension for visual purposes using a resolution of 2 Hz and were then thresholded to remove low power background.

### Fish identification and tracking

Our EOD tracking system is optimized for identifying and tracking individual wave-type electric fish, to estimate the fish’s positions, and to detect communication signals. The signals of pulse-type electric fish were detected, but remain unprocessed for now. First, information about electric fish presence, EOD frequency (EOD *f*), and approximate position were extracted. Each electrode signal was analyzed separately in sequential overlapping windows (1.22 s width, 85 % overlap). For each window the power spectral density was calculated (8192 FFT data points, 5 sub-windows, 50% overlap) and spectral peaks above a given threshold were detected. Individual fish were extracted from the list of peak frequencies, based on the harmonic structure of wave-type EODs. For each analysis window, EOD detections from all electrodes were matched and consolidated. Finally, fish detections in successive time windows were matched, combined, and stored for further analysis.

### Position estimation

Once the presence of an electric fish was established, the fish’s position was estimated and chirps were detected. For each fish, the signals of all electrodes were bandpass-filtered (forward-backward butterworth filter, 3rd order, 5*×* multipass, *±*7 Hz width) at the fish’s EOD *f*. Then the envelope was computed from the resulting filtered signal using a root-mean-square filter (10 EOD cycles width). Each 40 ms the fish position 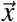 was estimated from the four electrodes *i* with the largest envelope amplitudes *A*_*i*_ at position 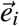 as a weighted spatial average

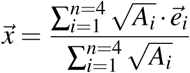

(movie M 1). If fewer than two electrodes with EOD amplitudes larger than 15 μV were available, the position estimate was omitted. Although this estimate does not relate to the underlying physics of the electric field, it proved to be the most robust against interference by electrical noise (Hopkins, 1973) and fish moving close to the edges of the electrode array, as verified with both experiments and simulations (Henninger, 2015). In short, we measured the spatial distribution of an electric fish’s EOD field in a large tank (3.5 *×* 7.5 *×* 1.5 m*, w × l × h*) under conditions similar to field conditions (water depth 60 cm, fish and electrode array submerged 30 cm below surface). We used this dataset for evaluating the performance of three algorithms for position estimation and for fitting a simple dipole model for the spatial electric field distribution. The dipole-model was then used to evaluate the algorithms in greater detail by simulating stationary and moving fish for various electrode configurations. For the electrode configuration used, the weighted spatial average yielded a precision of 4.2*±*2.6 cm on level with the electrode array and 6.2*±*3.8 cm at a vertical distance of 15 cm as computed by extensive simulations. Finally, the position estimates were filtered with a running average filter of 200 ms width to yield a smoother trace of movements. For pulse-type electric fish an EOD-based method for tracking electric fish position in shallow water in a laboratory setup has been published recently, yielding slightly better precision with a physically more realistic lookup-table-based approach (Jun et al., 2013).

### Chirp detection and analysis

For each fish the electrode voltage traces were bandpass-filtered (forward-backward butterworth filter, 3rd order, 5*×* multipass, *±*7 Hz width) at the fish’s EOD *f* and at 10 Hz above the EOD *f*. For each passband the signal envelope was estimated using a root-mean-square filter over 10 EOD cycles. Rapid positive EOD frequency excursions cause the signal envelope at the fish’s baseline frequency to drop and in the passband above the fish’s EOD *f* to increase in synchrony with the frequency excursion. If events were detected synchronously in both passbands on more than two electrodes, and exceeded a preset amplitude threshold, they were accepted as communication signals.

Communication signals with a single peak in the upper passband were detected as small chirps. Signals of up to 600 ms duration and two peaks in the upper passband, marking the beginning and the end of the longer frequency modulation, were detected as long chirps. All chirps in this study were verified manually. However, it is likely that some chirps were missed, since detection thresholds were set such that the number of false positives was very low. Also, abrupt frequency rises (AFRs, Engler and Zupanc, 2001) were probably not detected because of their low frequency increase.

Interchirp-interval probability densities were generated for pairs of fish and only for the time period in which both fish were producing chirps. Kernel density histograms of interchirp intervals (Fig. 5 — figure supplement 1) were computed with a Gaussian kernel with a standard deviation of 20 ms.

Rates of small chirps before and after female long chirps (Fig. 5 A, C) were calculated by convolving the chirp times with a Gaussian kernel (σ = 0.5 s) separately for each episode and subsequently calculating the means and standard deviations.

For quantifying the echo response (Fig. 6) we computed the cross-correlogram

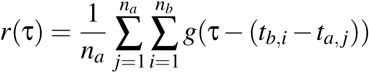

with the *n*_*a*_ chirp times *t*_*a,* *j*_ of fish *a* and the *n*_*b*_ chirp times *t*_*b,i*_ of fish *b* using a Gaussian kernel *g*(*t*) with a standard deviation of 20 ms. To estimate its confidence intervals, we repeatedly resampled the original dataset (2000 times jackknife bootstrapping; random sampling with replacement), calculated the cross-correlogram as described above and determined the 2.5 and 97.5 % percentiles. To create the cross-correlograms of independent chirps, we repeatedly (2000 times) calculated the cross-correlograms on chirps jittered in time by adding a random number drawn from a Gaussian distribution with a standard deviation of 500 ms and determined the mean and the 2.5 and 97.5 % percentiles. Deviations of the observed cross-correlogram beyond the confidence interval of the cross-correlogram of jittered chirp times are significant on a 5 % level, and are indicative of an echo response. Reasonable numbers of chirps for computing meaningful cross-correlograms (more than several hundreds of chirps) were available in five pairs of fish.

### Beat frequencies and spatial distances

The distance between two fish at the time of each chirp (Fig. 8 B) was determined from the estimated fish positions. Because position estimates were not always available for each time point we allowed for a tolerance of maximally two seconds around the chirp for retrieving the position estimate. For 95% of all chirps distance estimates were available within less than 400 ms. To evaluate the validity of our estimates in regard to fish movement occurring during the time of a chirp and the time of an estimate we calculated the the distance between the temporally closest available estimate before and after each chirp. We found that for 95% of all chirps the differences in distance estimates before and after the chirp were smaller 6.2 cm. The distance estimates were compiled into kernel density histograms that were normalized to their maximal value. The Gaussian kernel had a standard deviation of 1 cm for courtship small chirps, and 2 cm for courtship long chirps as well as intruder small chirps. Males (*n* = 8) intruding on a courting dyad initially lingered at some distance from the dyad before either approaching the dyad further or being chased away by the courting resident male. Distances between the intruding male and the courting male during this assessment behavior (Fig. 8 C, top) were measured every 40 ms beginning with the appearance of the intruding fish until the eventual approach or attack. These distances, collected from a total assessment time of 923 s, were summarized in a kernel density histogram with Gaussian kernels with a standard deviation of 2 cm.

When a male intruded on a courting dyad it was directly attacked by the resident male. In that process courtship was always interrupted and eventually one of the males withdrew. In some cases a few chirps were emitted by the retreating male. The winning male always approached and courted the female. The attack distances between two males (Fig. 8 C, bottom) were determined at the moment a resident male initiated its movement toward an intruding male. This moment was clearly identifiable as the onset of a linear movement of the resident male towards the intruder from plots showing the position of the fish as a function of time.

The distribution of beat frequencies generated by fish present in the electrode array at the same time (Fig. 8 E) was calculated from all available recordings during the breeding season. The average frequency difference of each pair of fish simultaneously detected in the recordings was compiled into a kernel density histogram with a Gaussian kernel with a standard deviation of 10 Hz. Similarly, for courtship and aggressive behavior (Fig. 8 F, G) the mean frequency differences were extracted for the duration of these interactions.

### Electric fields

For an estimation of EOD amplitude as a function of distance, histograms of envelope amplitudes from all electrodes of the array were computed as a function of distance between the electrodes and the estimated fish position. For each distance bin in the range of 20 – 100 cm the upper 95 % percentile of the histogram was determined and a power law was fitted to these data points. Gymnotiform electroreceptors measure the electric field, i.e., the first spatial derivative of the EOD amplitudes as shown in Fig. 8 A.

### Breeding monitoring setup

In the laboratory breeding study, we used the brown ghost knifefish *Apteronotus leptorhynchus*, a close relative of *A. rostratus* (de Santana and Vari, 2013). *Apteronotus leptorhynchus* is an established model organism in neuroscience and readily available from aquarium suppliers. The two species share many similarities. (i) Most chirps produced by both species are “small chirps” that in *A. leptorhynchus* have been classified as type-2 chirps (Engler and Zupanc, 2001). (ii) Females of both species additionally generate small proportions of “long chirps”, similar to the type-4 chirps classified for *A. leptorhynchus* males. (iii) Both species show the same sexual dimorphism in EOD *f*.

The laboratory setup for breeding *A. leptorhynchus* consisted of a tank (100 *×* 45 *×* 60 cm) placed in a darkened room and equipped with bubble filters and PVC tubes provided for shelter. Heating was supplied via room temperature, which kept water temperature between 21 and 30°C. The light/dark cycle was set to 12/12 hours. Several pieces of rock were placed in the center of the tank as spawning substrate. EOD signals were recorded differentially using four pairs of graphite electrodes. Two electrode pairs were placed on each side of the spawning substrate. The signals were amplified and analog filtered using a custom-built amplifier (100*×* gain, 100 Hz high-pass, 10 kHz low-pass; npi electronics GmbH, Tamm, Germany), digitized at 20 kHz with 16 bit (PCI-6229, National Instruments, Austin, Texas, USA), and saved to hard disk for offline analysis. The electrode pairs were positioned orthogonally to each other, thereby allowing for robust recordings of EODs independent of fish orientation. The tank was illuminated at night with a dozen infrared LED spotlights (850 nm, 6W, ABUS TV6700) and monitored continuously (movie M 4) with two infrared-sensitive high-resolution video cameras (Logitech HD webcam C310, IR filter removed manually). The cameras were controlled with custom written software (https://github.com/bendalab/videoRecorder) and a timestamp for each frame was saved for later synchronization of the cameras and EOD recordings. Six fish of *A. leptorhynchus* (three male, three female; imported from the Rίo Meta region, Colombia) were kept in a tank for over a year before being transferred to the recording tank. First, fish were monitored for about a month without external interference. We then induced breeding conditions (Kirschbaum and Schugardt, 2002) by slowly lowering water conductivity from 830 μS*/*cm to about 100 μS*/*cm over the course of three months by diluting continuously the tank water with deionized water. The tank was monitored regularly for the occurrence of spawned eggs.

### Electrophysiology

For *in vivo* recordings fish were anesthetized with MS-222 (120 mg/l; PharmaQ, Fordingbridge, UK; buffered to pH 7 with sodium bicarbonate) and a small part of the skin was removed to expose the anterior part of the lateral line nerve that contains only electroreceptor afferent fibers innervating electroreceptors on the fish’s trunk (Maler et al., 1974). The margin of the wound was treated with the local anesthetic Lidocaine (2%; bela-pharm, Vechta, Germany). Then the fish was immobilized by intramuscular injection of Tubocurarine (Sigma-Aldrich, Steinheim, Germany; 25 – 50 μl of 5 mg/ml solution), placed in a tank, and respired by a constant flow of water through its mouth. The water in the experimental tank (47 *×* 42 *×* 12 cm) was from the fish’s home tank with a conductivity of about 300 μS*/*cm and kept at 28 °C. All experimental protocols were approved by the local district animal care committee and complied with federal and state laws of Germany (file no. ZP 1/13) and Canada.

Population activity in whole-nerve recordings was measured using a pair of hook electrodes of chlorided silver wire. Signals were differentially amplified (gain 10 000) and band-pass filtered (3 to 3 000 Hz passband, DPA2-FX; npi electronics), digitized (20 kHz sampling rate, 16-bit, NI PCI-6229; National Instruments), and recorded with RELACS (www.relacs.net) using efield and efish plugins. The strong EOD artifact in this kind of recording was eliminated before further analysis by applying a running average of the size of one EOD period (Benda et al., 2006). The resulting signal roughly followed the amplitude modulation of the EOD and we quantified its amplitude by taking its standard deviation. The nerve recordings closely resemble the properties of P-unit responses obtained from single and dual-unit recordings (Benda et al., 2006; Walz et al., 2014). Note, however, that P-units might still respond in subtle ways to a stimulus even though the nerve recording is already down at baseline level, because of additional noise sources in this kind of recording. Also note, that the lateral-line nerve only contains fibers from one side of the fish’s body.

Electric sine-wave stimuli with frequencies ranging from Δ *f* = *-*460 to +460 Hz in steps of 2 Hz (*|*Δ *f | ≤* 20 Hz), 10 Hz (*|*Δ *f | ≤* 200 Hz), and 20 Hz (*|*Δ *f | >* 200 Hz) relative to the fish’s EOD *f* were applied through a pair of stimulation electrodes (carbon rods, 30 cm long, 8 mm diameter) placed on either side of the fish. Stimuli were computer-generated and passed to the stimulation electrodes after being attenuated to the right amplitude (0.05, 0.1, 0.2, 0.5, 1.0, 2.5, 5.0, 10.0, 20.0, 40.0 % of the fish’s EOD amplitude estimated with a pair of electrodes separated by 1 cm perpendicular to the side of the fish) and isolated from ground (Attenuator: ATN-01M; Isolator: ISO-02V; npi electronics). For more details see Benda et al. (2006) and Walz et al. (2014). Using Fig. 8 A the stimulus amplitudes were placed on the corresponding distances.

## Acknowledgment

Supported by the BMBF Bernstein Award Computational Neuroscience 01GQ0802 to J.B., a Discovery Grant from the Natural Sciences and Engineering Research Council of Canada to RK, and a Short Time Fellowship to J.H. from the Smithonian Tropical Research Institute. We thank Hans Reiner Polder and Jürgen Planck from npi electronic GmbH for designing the amplifier, Sophie Picq, Diana Sharpe, Luis de León Reyna, Rigoberto González, Eldredge Bermingham, the staff from the Smithsonian Tropical Research Institute, and the Emberá community of Peña Bijagual for their logistical support, Fabian Sinz for advice on the analysis, and Ulrich Schnitzler and Janez Presern for comments on the manuscript.

## Competing interests

Authors declare no conflicts of interest.

## Audio

### Animations and Video

**Audio** A 1: Audio trace of the courtship sequence shown in Fig. 4. A male (EOD *f* = 930 Hz) generated a series of small chirps. Eventually, the female (EOD *f* = 620 Hz) fish joins in, increases chirp rate and finishes with a long chirp, which is acknowledged by the male with a small chirp doublet.

File: audio courtship.wav

**Movie** M 1: Example of raw voltage recordings and corresponding position estimates of a single fish, *Eigenmannia humboldtii*, passing through the array of electrodes. The head and tail area of its electric field are of opposite polarity, which is why the polarity of the recorded EOD switches as the fish passes an electrode. Note the large electric spikes occurring irregularly on all electrodes. Previous studies (Hopkins, 1973) attributed similar patterns to propagating distant lightning. The animation is played back at real-time.

File: movie raw and position.avi

**Movie** M 2: Animation of the courtship and aggression behavior shown in Fig. 2. A courting dyad is engaged in intense chirp activity (transparent circles and 50 ms beeps at the fish’s baseline EOD *f*). An intruder male (red circles indicate positions of the last 5 seconds, black circles mark current positions) first lingers at a distance of one meter. When it approaches further, courting is interrupted and the resident male engages the intruder. Just before the male intruder retreats, it emits a series of small chirps, and subsequently leaves the recording area. The resident male returns to the female and resumes chirping. Eventually, the female responds with small chirps followed by a single long chirp (large open circle and a 500 ms beep at the female’s baseline EOD *f*). Then both fish cease chirp activity and the male resumes to emit chirps after a few seconds. The animation is played back at 2*×* real-time.

File: movie intruder.avi

**Movie** M 3: Animation of a courtship sequence with multiple attempts of an intruding male to approach the courting dyad. The resident male drives the intruder away three times, starting the approach at increasingly greater distances. *Apteronotus rostratus* are marked by circles, *Eigenmannia humboldtii* by squares. The animation is played back at 2*×* real-time.

File: movie repetitive intruder.avi

**Movie** M 4: Spawning of the closely related species *Apteronotus lepthorhynchus* during a breeding experiment. The overall sequence of chirp production is very similar to the courtship motif observed in *A. rostratus*. However, male *A. lepthorhynchus* increasingly generate a second type of chirp, a variety of a long chirp, as spawning approaches. The video shows a big male (EOD *f* = 770 Hz) courting a smaller female (590 Hz). The audio signal was created from concurrent EOD recordings. Both fish generate chirps at an increased rate (about 1.5 Hz), just before the male thrusts its snout against the female, which responds with a long chirp, clearly noticeable from the audio trace. Subsequently, the male retreats to a tube and the female hovers around the substrate, where the spawned egg was found.

File: movie spawning.avi

## Figure supplements

**Figure 1 — figure supplement 1:**
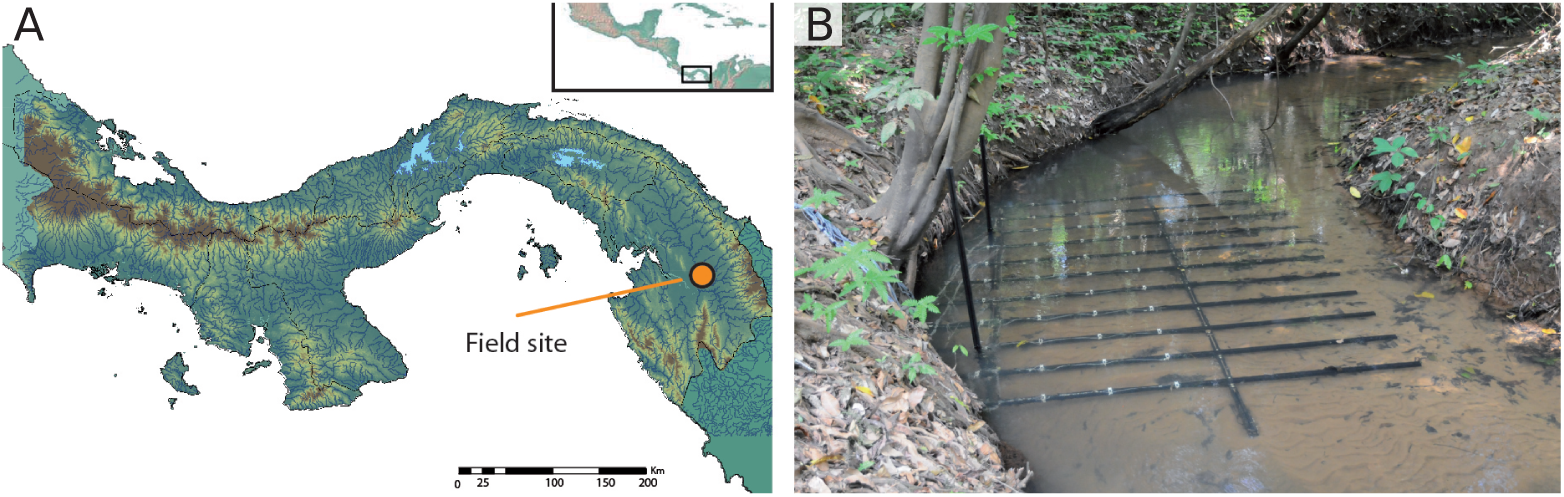
Field site and the electrode array positioned in a stream. A) The field data were recorded in the Darién province in Eastern Panamá. B) The electrode array covered 2.4 *×* 1.5 m^2^ of our recording site in a small quebrada of the Chucunaque River system. Electrodes (on white electrode holders) were positioned partly beneath the excavated banks, allowing to record electric fish hiding deep in the root masses.

**Figure 5 — figure supplement 1:**
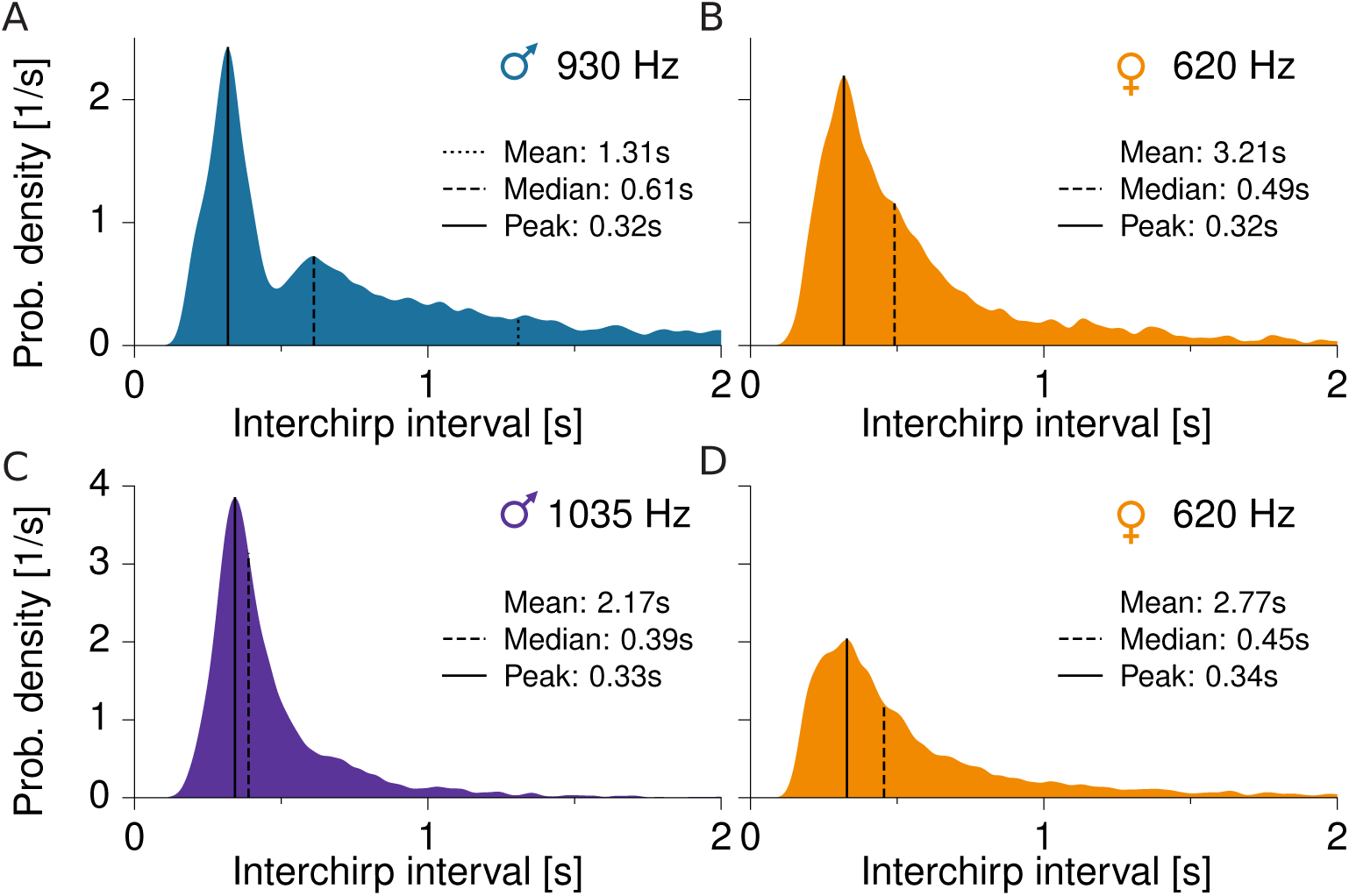
Interchirp-interval distributions of small chirps underlying the chirprates shown in fig. 5. A) Male with EOD *f* = 930 Hz (*n* = 8439 small chirps). B) Female with EOD *f* = 620 Hz (*n* = 3431). C) Another male with EOD *f* = 1035 Hz (*n* = 6857). D) Same female as in panel B (*n* = 5336 chirps).

**Figure 5 — figure supplement 2:**
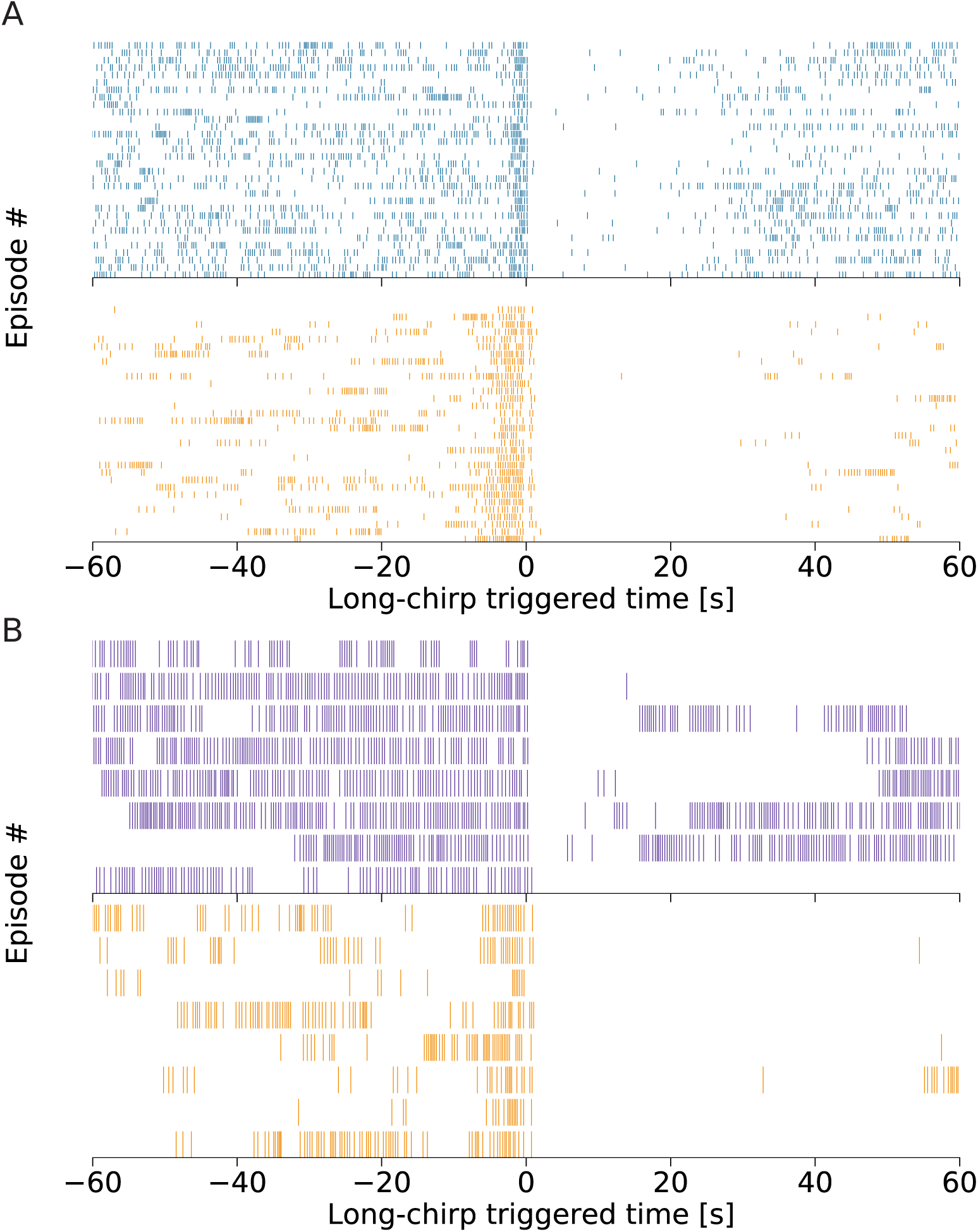
Raster plots of small chirps underlying the chirprates shown in fig. 5. A) Male with EOD *f* = 930 Hz (top) and female with EOD *f* = 620 Hz (bottom). B) Another male with EOD *f* = 1035 Hz (top) and same female as in panel A (bottom). Each row corresponds to a single courtship episode, each stroke marks a small chirp.

